# On a mechanistic impact of transmembrane tetramerization in pathological activation of RTKs

**DOI:** 10.1101/2022.10.14.512270

**Authors:** Anton A. Polyansky, Roman G. Efremov

## Abstract

Constitutive activation of receptor tyrosine kinases (RTKs) via different mutation has a strong impact into development of severe human disorders, including cancer. While pathological effect of such mutations can be common on the phenotypical level, mechanistic understanding of their contribution into the receptor activation depends on the exact molecular context. Mutations in transmembrane (TM) domains represent an interesting class of pathological modifications since they can directly affect a signal transduction pathway from the receptor to the kinase domains of RTKs. Here we propose a putative activation scenario of RTK, whereby TM mutations can also promote higher order oligomerization of the receptors that leads to the subsequent ligand-free activation. To illustrate this model with all-atom resolution for a previously characterized oncogenic TM mutation V536E in platelet-derived growth factor receptor alpha (PDGFRA), we use a computational modeling framework including sequence-based structure prediction and all-atom 1 µs molecular dynamics (MD) simulations in a model membrane for the predicted configuration of the PDGFRA TM tetramers. We show that in the course of MD simulations the mutant tetramer retains stable and compact configuration, which is strengthened by tight protein-protein interactions. In contrast, the wild type tetramer demonstrates looser packing and tendency to dissociate. Such a structural organization shapes the dynamics of TM helices in the oligomeric state. Specifically, the mutation affects the characteristic motions of mutated TM helical segments by introducing additional non-covalent crosslinks in the middle of the TM tetramer, which work as mechanical hinges. This leads to dynamic decoupling of the C-termini from the rigidified N-terminal parts and facilitates higher possible displacement between the C-termini of the mutant TM helical regions. This, in turn, can provide more freedom to downstream kinase domains in their mutual rearrangement. The observed structural and dynamic effects of the V536E mutations in the context of PDGFRA TM tetramer provide an interesting possibility that an effect of oncogenic TM mutations can go beyond alternating structure and dynamics of TM dimeric states and also promote formation of higher-order oligomers that may directly contribute into the ligand independent signaling of PDGFRA and other RTKs.

## 1. Introduction

Receptor-tyrosine kinases (RTKs) undergo dimerization or high order oligomerization upon functioning (M. D. Paul & K. Hristova, 2019) (Westerfield & Barrera, 2020). While RTK dimerization is typically in focus of biochemical and biophysical studies, the latter aspect and its direct impact into receptor activation is far from being fully understood especially in the context of oncogeneses and other functional disorders. For instance, platelet-derived growth factor receptor alpha (PDGFRA), is aberrantly activated in many neoplasms (e.g., glioblastoma), where its pathologic activation is caused by gene amplification, gene fusion, autocrine production of the ligand or point mutations (Andrae, Gallini, & Betsholtz, 2008) (Andrae et al., 2008) (Demoulin & Essaghir, 2014). Thus, we have previously shown that V536E substitution in the PDGFRA transmembrane (TM) domain is sufficient to constitutively activate the receptor (Velghe et al., 2014). Moreover, for isolated TM domains of V536E variant it was experimentally observed a prominent tendency to form oligomers of higher order than dimers. The latter was confirmed by SDS page and characteristic NMR spectra for different protein-to-lipid ratio (A. A. Polyansky et al., 2019). At the same time, the oligomerization was impaired for TM of the WT protein, where only moderate dimerization tendency was detected by the aforementioned methods. One reasonable hypothesis here can be that the oncogenic V536E mutation potentially modulates the receptor activity by shifting the equilibrium toward formation of higher oligomers that subsequently facilitates ligand-free activation by increasing the probability of autophosphorylation of kinase domains. First, oligomers promote higher local concertation of the catalytic domains. Second, oligomers geometry and dynamics can potentiate the asymmetric arrangement and association of the kinase domains even in the absence of the signal. Thus, basal activation of the relative PDGFRB receptor can be achieved via TM hetero-oligomerization in the presence of E5 oncoprotein from bovine papillomavirus (Karabadzhak et al., 2017). Also, for PDGFRB it was shown integrin-mediated constitutive activation (Sundberg & Rubin, 1996) (Seong, Huang, Sim, Kim, & Wang, 2017), whereby β1-integrin and PDGFRB clustering associated with tyrosine phosphorylation of the receptor (Sundberg & Rubin, 1996). While up to date the possibility to form higher-order oligomers for PDGFRA receptor is rather under debate, such a trend was observed for another RTKs, e.g. for ephrin receptors (Michael D. Paul & Kalina Hristova, 2019) or epidermal growth factor receptor (EGFR). Thus, ligand-free and ligand-induced oligomerization of EGFR was connected to autoinhibition (Zanetti-Domingues et al., 2018) or fully-activate state (Clayton et al., 2005) (Clayton, Orchard, Nice, Posner, & Burgess, 2008) of the receptor and formation of signaling platforms with enhanced (‘superstoichiometric’) response to EGF stimulation (Needham et al., 2016), respectively.

Importantly to note that not only RTKs but many others membrane proteins realize their function by direct binding to each other and interacting via different domains including the TM ones. That is why mechanistic understanding of oligomerization of TM helices represents a topic of a general interest. The latter requires quite laborious experiential studies complicated also by highly heterogenous membrane environment that hard to mimic *in vitro* and to control *in vivo*. Likely, recent advances in biophysical techniques based on Förster resonance energy transfer enable *in vivo* studies of signaling and dynamics of membrane protein association specific to different membrane microdomains using specific fluorescent biosensors (Zhang, Mehta, & Zhang, 2021). Such experiments highlight an importance of receptor localization in specific membrane environment and/or oligomerization for the signal transduction through the membrane. For instance, already in dimeric form TM domains (particularly, in the case of RTKs) gain novel characteristics, which are absent at the monomeric stage (e.g., (A. A. Polyansky, Volynsky, & Efremov, 2012)). Thus, TM dimers can adopt different functional states and transitions between them usually underly the protein functioning (see (Kovacs, Zakany, & Nagy, 2022) for a recent review). Although a number of studies in the last decade are focused on structure and dynamics of TM dimers and corresponding protein functions (e.g., active/inactive RTK dimers (Endres et al., 2013) (Bocharov, Bragin, et al., 2017)), many questions remain still open. For instance, in the case of RTKs these are mapping of exact allosteric pathways and contribution of characteristic dynamics of TM helices into the signal transduction between receptor and kinase domains. Even much less is known about dynamics and structural organization of higher order TM oligomers, e. g., tetramers. In the latter case a key question in analogy to the dimers: does a tetramer resemble just a simple sum of its composing parts (TM monomers and dimers) or does tetramerization due to synergy render novel properties of the system, which are absent at the level of monomers and dimers? Answering this fundamental question is crucial for understanding the molecular background of cellular signaling, etc., whereby atomistic modeling in combination with current biophysical techniques provide an efficient research framework as was previously shown for PDGFRA (A. A. Polyansky et al., 2019).

In the present study we use different modeling techniques to study effect of V536E mutation in the context of TM tetramers in comparison to the WT. To achieve this purpose, we reconstruct tetramer architecture for the both variants using assembly of dimers predicted by PREDDIMER approach (Anton A. Polyansky et al., 2013) (A. A. Polyansky et al., 2019), and explore their structural and dynamic organization in the model lipid membrane using all-atom 1 µs molecular dynamics (MD) simulations.

## 2. Methods

### 2.1 Reconstruction of tetrameric configuration for PDGFRA TM

TM sequence fragment ^522^-SELTVAAAVLVLLVIVIISLIVLVVIWKQ-^551^ of human PDGFRA (Uniprot ID: P16234) was used for WT and V536E mutant, whereby the corresponding substitution (V->E) was applied at 536^th^ position. Left-handed (L_I_^N^) and right-handed (R_II_^C^) TM dimer configuration (Figure 1A) were obtained using PREDDIMER web-server (Anton A. Polyansky et al., 2013) and combined into a tetrameric arrangement as described also elsewhere (A. A. Polyansky et al., 2019).

**Figure 1.**
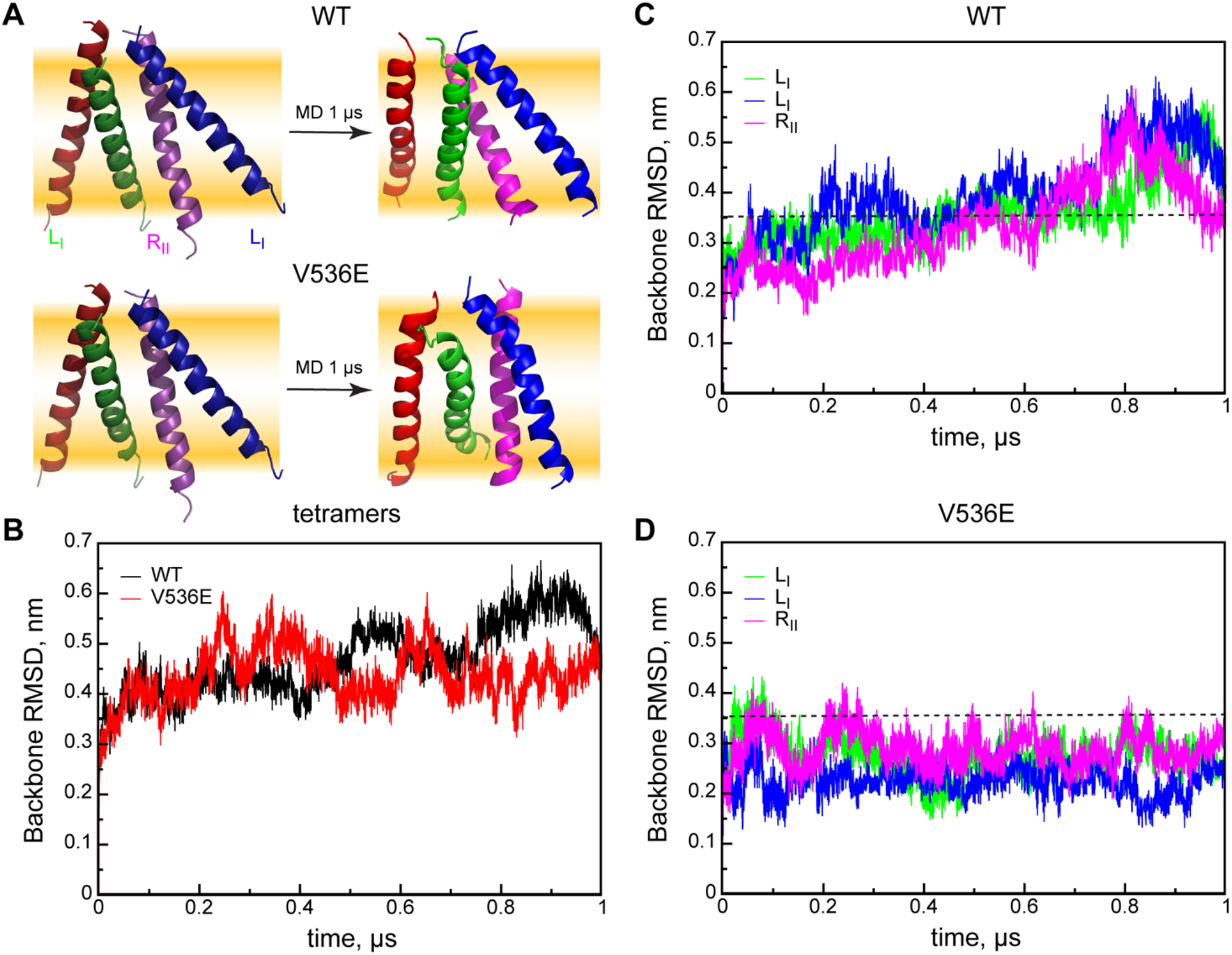
Evolution of tetrameric configurations during 1 µs MD. **A**. Starting (*left*) and final (*right*) conformations of WT (*up*) and V536E (*bottom*) tetramers from the side view. TM helices are shown with cartoon representation and colored as follows: red and green – the first left-handed dimer (L_I_); magenta and blue – the second left-handed dimer (L_II_). The central right-handed dimer (R_ii_) is formed by green and magenta helices. POPC lipid bilayer is schematically shown with orange gradients. **B**. Backbone root-mean-square deviations (RMSD) from the start of the tetrameric conformation during 1 µs MD simulations for the WT (black) and V536E (red). **C**. Backbone RMSDs from the start for individual dimers within the WT tetramer structure. The first and the second left-handed dimers are shown with green and blue curves, respectively. The right-handed dimer is shown with a magenta curve. RSMD value of 0.5 nm indicated with a dashed line. **D**. Backbone RMSDs from the start for individual dimers within the V536E tetramer structure. The color code is the same as in panel C.

### 2.2 MD simulations

All MD simulations were performed using GROMACS 4.5 package (Pronk et al., 2013) and Amber99SB-ILDN force-field (Lindorff-Larsen et al., 2010) with lipid parameters (Slipids) (Jambeck & Lyubartsev, 2012) and TIP3P water model (Jorgensen, 1981) using the similar protocol as described elsewhere (Jambeck & Lyubartsev, 2012) (A. A. Polyansky et al., 2019). Briefly, the initial configurations of the WT and V536E systems were obtained using *genbox* from the GROMACS package by inserting reconstructed tetramer conformations into a pre-equilibrated 1-palmitoyl-2-oleoyl-sn-glycero-3-phosphocholine (POPC) lipid bilayer comprised of 200 molecules (7×7×12 nm^3^ box). The final systems each contain 1 protein, 129 lipids, ∼14 10^4^ water molecules. These systems were energy minimized and subjected to a few steps MD equilibration with positionally constrained protein heavy atoms in X- and Y-directions. Equilibration steps were performed in different thermodynamic ensembles and using gradually increasing integration time step: 5000 steps /0.5 fs / NVT, 25000 steps / 0.5 fs / NPT semi-isotropic pressure (Berendsen barostat (Berendsen, Postma, van Gunsteren, DiNola, & Haak, 1984)), 250000 / 1 fs / NPT (Parrinello-Rahman barostat (Parrinello & Rahman, 1981)). Finally, production runs of 1 µs were carried out for the both systems using a 2 fs time step. A twin-range (0.10/0.14 nm) spherical cut-off function was used to truncate van der Waals interactions. Electrostatic interactions were treated using the particle-mesh Ewald summation (real space cutoff 1.0 and 0.12 nm grid with fourth-order spline interpolation). 1µs MD simulations were carried out using 3D periodic boundary conditions in the isothermal−isobaric (NPT) ensemble with an semisotropic pressure of 1.013 bar for both directions and a constant temperature of 310K. The pressure and temperature were controlled during the 1µs production runs using Nose-Hoover thermostat (Hoover, 1985) and a Parrinello-Rahman barostat (Parrinello & Rahman, 1981) with 0.5 and 10 ps relaxation parameters, respectively, and a compressibility of 4.5 × 10^−5^ bar^−1^ for the barostat for both directions. During the simulations the box sizes were adjusting and equilibrating around the following dimensions: to 6.7×6.7×13.2 nm^3^. Protein, lipids and water molecules were coupled separately. Bond lengths were constrained using LINCS (Hess, Bekker, Berendsen, & Fraaije, 1997).

### 2.3 MD analysis

All analyses were done using utilities from the GROMACS 5 package (Abraham et al., 2015) and Linux scripts specially written for this purpose. The following MD-derived parameters were calculated using GROMACS utilities: root-mean-square deviation from the starting configuration and a reference MD trajectory (RMSD, *rms*), radius of gyration (*Rg, gyrate*), MD distances between pairs of atoms (*pairdist*). A number of van-der-Waals contacts between protein residues was calculated with an applied 0.35 nm cut-off for any interatomic distances between the two moieties. Clustering of MD tetramer conformations was performed using backbone RMSD cut-off defining the cluster perimeter from 0.05 to 0.15 nm with a 0.02 nm step (Figure S1). An optimal cut-off value of 0.1 nm was selected for further analysis. The characteristic tetramer dynamics were studied using calculations of eigenvectors from mass-weighted variance-covariance matrices obtained for MD distribution of x-,y-,z-atomic degrees of freedom (*covar*). 1µs MD trajectories were filtered (*anaeig*) along the first three eigenvectors (1-3, *anaeig*) and further used for visualization of dynamics of TM helices and calculations of atomic fluctuations (root-mean-squared fluctuation, RMSF, *rmsf*) corresponding to these components. All structures were visualized using Pymol (Schrodinger, 2010).

## 3. Results

### 3.1 PDGFRA TM domains are able to form a tetrameric configuration

Analysis of different dimer conformations possible for PFGRA TM revealed the two dimerization interfaces located on the opposite sides of the helix (A. A. Polyansky et al., 2019). While the left-handed dimers for the mutant and WT have very similar configuration, in which helices are associated via interface I of the N-terminal parts (it contains the mutated V536, LI^N^-dimer), the right-handed dimers are different in these two cases. Thus, a unique feature of the mutant TM is the possibility to form relatively stable right-handed dimer, where helices associate along interface II in the C-terminal region (R_II_^C^-dimer). The latter gives a possibility for the mutant TM helices to attend a tetrameric configuration, where the R_II_^C^-dimer represents a core, while additional helices associate along the interface I engaging E536 for inter-helical contacts and resulting in the formation of two additional L_I_^N^-dimers. Such a configuration fits to NMR data obtained for the V536E mutant at lower detergent-to-protein ratio (A. A. Polyansky et al., 2019). Thus, both the WT and the V536E mutant TM-tetramers were built in the described configuration (**Figure 1A**) and subjected to all-atom MD simulations in a POPC lipid bilayer (see below). Interestingly, L_I_^N^-dimer represents a conservative TM configuration and was found also for PDGFRB (Muhle-Goll et al., 2012). A potential scenario for the receptor tetramerization can be that at first it forms dimers where TM helices adopt L_I_^N^-configurations and later these dimers interact subsequently with each other and form additional R_II_^C^-dimer at the interface. Similar scenario albeit without higher-order oligomerization in TM was proposed for tetramerization of EGFR (Needham et al., 2016).

### 3.2 V536E forms stable and converged tetramer configuration in MD

During MD simulations both tetramers display rearrangement of the initial configuration that is reflected in the root-mean-square deviation (RMSD) from the start calculated for the backbone atoms (**Figure 1B**). Thus, RMSD for the WT tetramer is gradually increasing up to 0.6 nm and then decreases to 0.5 nm at the very end of the simulation. In contrast, evolution of V536E tetramer resembles forward-and-back character and for most parts of the trajectory remains below 0.5. In the latter case, the tetramer preserves in general the initial configuration (**Figure 1A, D**), whereby two L_I_^N^ dimers and the central R_II_^C^ dimer display RMSD below 0.35 nm during the MD trajectory. In contrast, in the WT tetramers helices do not preserve stable arrangement. For instance, one of L_I_^N^ dimers (blue/magenta helices, **Figure 1A, D**) in WT tends to unfold, whereby helices loose contacts in the N-terminal region and the first helix (blue) forms a new N-terminal dimer with another partner, while the second one (magenta) retains only contacts with other helices in the C-terminal part (**Figure 1A**). At the same time, the V536E mutation stabilizes the left-handed dimers along the interface I and facilitates formation of stable R_II_^C^ dimer that directly contributes into the stability and the structural organization of the mutant tetramer. Detailed structural cluster analysis of tetrameric configurations (**Figure S1**) demonstrates a convergence behavior for the mutant, whereby its conformation is gradually evolving (**Figure S1B**) toward the dominating state at the last part of MD trajectory shown in **Figure S1C** (a representative structure for the most occupied structural cluster 1 with the occupancy of 17.2 %). The convergence of the tetrameric configuration is highlighted in decreasing of the mutual RMSD values between the representative structures from three top clusters (**Figure S1D**). In contrast, the WT tetramer forms the dominating conformation (cluster 1, 16%) in the middle of the MD trajectory, which evolved further into the less populated state (cluster 2, 9.8%) unstable toward the end of the simulations. Such an evolution is characterized by gradually increasing RMSD values between representative conformations of three top clusters along the MD time (**Figure S1D**). Importantly, the mutant tetramer remains a tight configuration during MD with a trend to further compaction (**Figure 2A, B**), while WT demonstrates opening of the helical bundle and accommodation of lipid molecules in the tetramer interior (**Figure 2A, B**). Thus, the number of protein-protein contacts remains stable for the mutant tetramer during MD and is almost halved for the WT (**Figure 2C**). The latter clearly indicates a transient nature of the WT tetramer with a tendency to dissociation. Moreover, compact organization of the mutant bundle with well-preserved configuration of both L_I_^N^ dimers facilitates wider lateral separations between the C-termini as compared to WT (**Figure 2D**).

**Figure 2.**
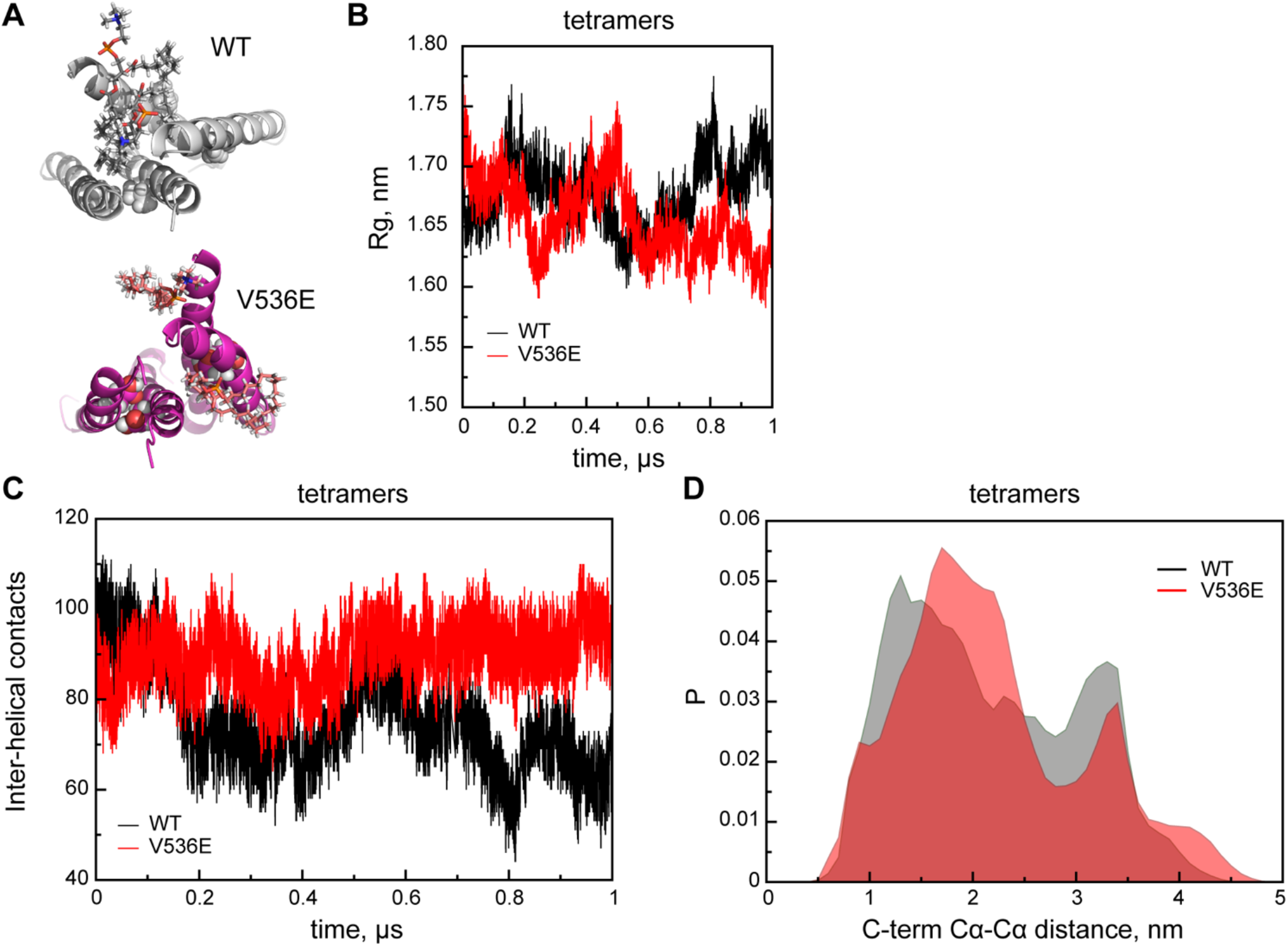
Tetramer packing and mutual arrangement. **A**. Final snapshots of the WT (grey color) and the V536E (magenta color) tetramers from the top view shown in cartoon representation with the closest lipid neighbors shown with sticks. **B**. MD evolution of the radius of gyration (*Rg*) calculated for tetramer conformations of the WT (black) and V536E (red). **C**. A total number of inter-helical atomic protein contacts for the WT (black) and V536E (red) tetramers. **D**. MD distributions of pairwise distances between Cα-atoms of the C-terminal residues in the WT (dark grey) and V536E (red) tetramers.

### 3.3 V536E mutation decouples dynamics of N- and C-termini within the tetramer

Since the transmission of allosteric signal between receptor and kinase parts of RTKs assumes a characteristic pattern in dynamics of TM helices, whereby different perturbation in its N-terminal parts is directly transmitted towards the C-termini, dynamic effect of the mutation was investigated in the tetramer context. The principal component analysis of MD-derived covariance matrices demonstrates a prominent difference between dynamics in the WT and the mutant tetramer. While only the first 10 obtained eigenvalues are substantially different between the two systems, the first 3 eigenmodes give about 60 % coverage of the statistical variation in WT and less in V536E – 50 %. (**Figure 3A**). Analysis of coordinate variances during MD associated with the corresponding eigenvectors (or RMSF values for MD trajectories filtered along the first 3 eigenvectors) demonstrates clearly different dynamic pattern along TM helices in WT and mutant tetramer (**Figure 3B**). In this case, WT helices accumulate most of dynamics in the N-terminal parts, while this effect is shifted to the C-termini in the mutant. Thus, in the mutant tetramer the C-termini display highest mobility, while the rest of the tetramer is relatively rigid (**Figure 3C**). Dynamic difference of the WT and the mutant is clearly seen in visualization of MD trajectories filtered along the first 3 principal components (**Movie S1, S2**). WT helices demonstrate piston-like movements en block with a tendency of the bundle to fall apart (**Movie S1**). At the same time, the mutant helices demonstrate motions with smaller amplitude, whereby most of dynamics is located in the C-terminal part (**Movie S2**). Interestingly that the mutant helices loose translational freedom and look apparently glued in the middle part by polar Glu sidechains.

**Figure 3.**
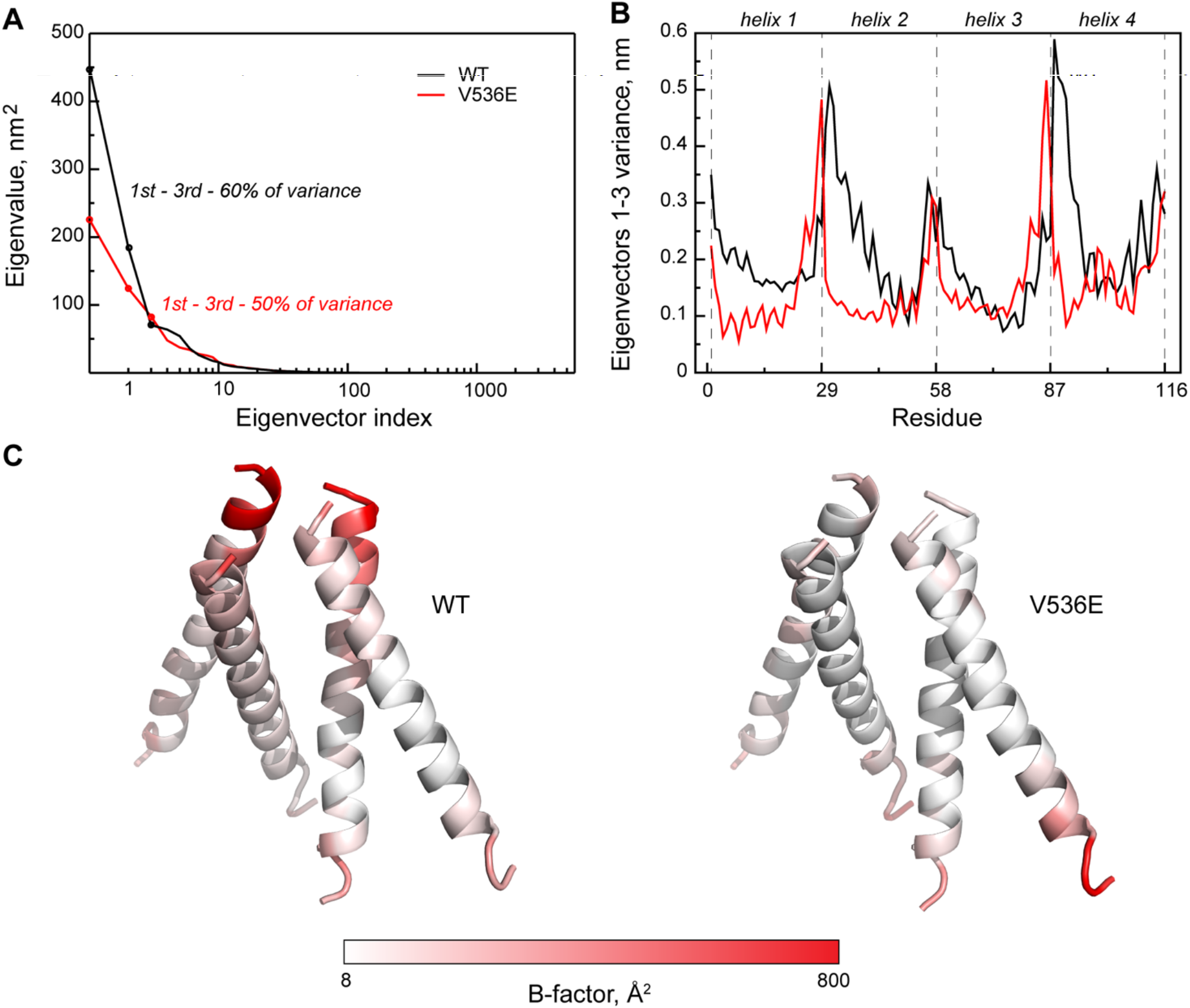
Characteristic dynamics of TM helices in tetrameric configuration. **A**. Distributions of eigenvalues for the mass-weighted MD covariance matrix for WT (black) and V536E (red) tetramers. First three eigenvalues are shown with filled circles. **B**. Sequence profiles of per residue atomic root-mean-square fluctuations (RMSF) along the first three eigenvectors for WT (black) and V536E (red) tetramers. Regions corresponding to individual TM helices are shown with vertical dashed lines. **C**. 3D distribution of atomic RMSFs along the first three eigenvectors in tetramer structures of the WT (*left*) and V536E (*right*). Tetramers are shown in cartoon representation. Helices are colored according to the corresponding B-factor scale given below the panel.

## 4. Discussion

The results of the modeling of two variants of the PDGFRA TM tetramer give a possibility to provide potential mechanistic explanations of the constitutive receptor activation via oligomerization induced by the oncogenic mutation, which are summarized in **Figure 4**. In the case of WT, TM helices are able to form a dimeric configuration (presumably, L_I_^N^-dimer) that corresponds to inactive state of the receptor as suggested before (A. A. Polyansky et al., 2019), while the further oligomerization step is impaired due to the observed transient and unstable nature of the tetrameric state. That is why, in the absence of the ligand (PDGF), WT PDGFRA most likely resides in inactive dimeric states whereby its activation can be achieved only upon the ligand binding and allosterically induced rearrangement of TM helices into active configuration. Importantly, that in the WT dimers (A. A. Polyansky et al., 2019) and the WT tetramer the helices demonstrate translational piston-like motions and directly coupled disposition of their N- and C-terminal parts suggesting that the mutual arrangement of the C-termini can be modulated only simultaneously with those of the N-termini (see **Movie S1**). The introduction of polar Glu residue in the middle of the PDGFRA TM helix as a result of the phenotypic mutation affects both the oligomerization of the mutant TM domains and their characteristic dynamics. As it was shown, the V536E mutants are able to form stable and well-defined tetrameric state, which in its turn may represent a nucleation step for subsequent oligomerization. Indeed, pre-formed mutant tetramers can further associate along the exposed R_Ii_ interfaces thus initiating a chain process, which would be limited by local concertation of the receptor. The latter process leads to high concertation of the kinase domains in local proximities, which by itself can increase the chance of spontaneous formation of their active configurations. The oligomerization potential of the isolated V536E TM domains was previously observed experimentally (A. A. Polyansky et al., 2019), with a trend to form discrete tetrameric state and the next iteration of suggested oligomerization pathway (e.g., octamers). Moreover, similar to the dimer (A. A. Polyansky et al., 2019) the mutant TM tetramer – a “building block” in high-order oligomerization – demonstrates alternated dynamic pattern assuming decoupling between the N- and C-termini due to the presence of non-covalent Glu crosslinks in the middle part. Namely, changes in configuration of the C-terminal parts can occur independently of that for the N-terminal ones in the absence of any allosteric impulse (see **Movie S2**). As it was shown, the C-terminal parts indeed are the most dynamic regions in the tetramer and display large mutual displacement that in context of the full-length receptor would give more freedom for downstream kinase domains to potential rearrangement into asymmetric configurations initiating autophosphorylation and subsequent signaling cascade.

**Figure 4.**
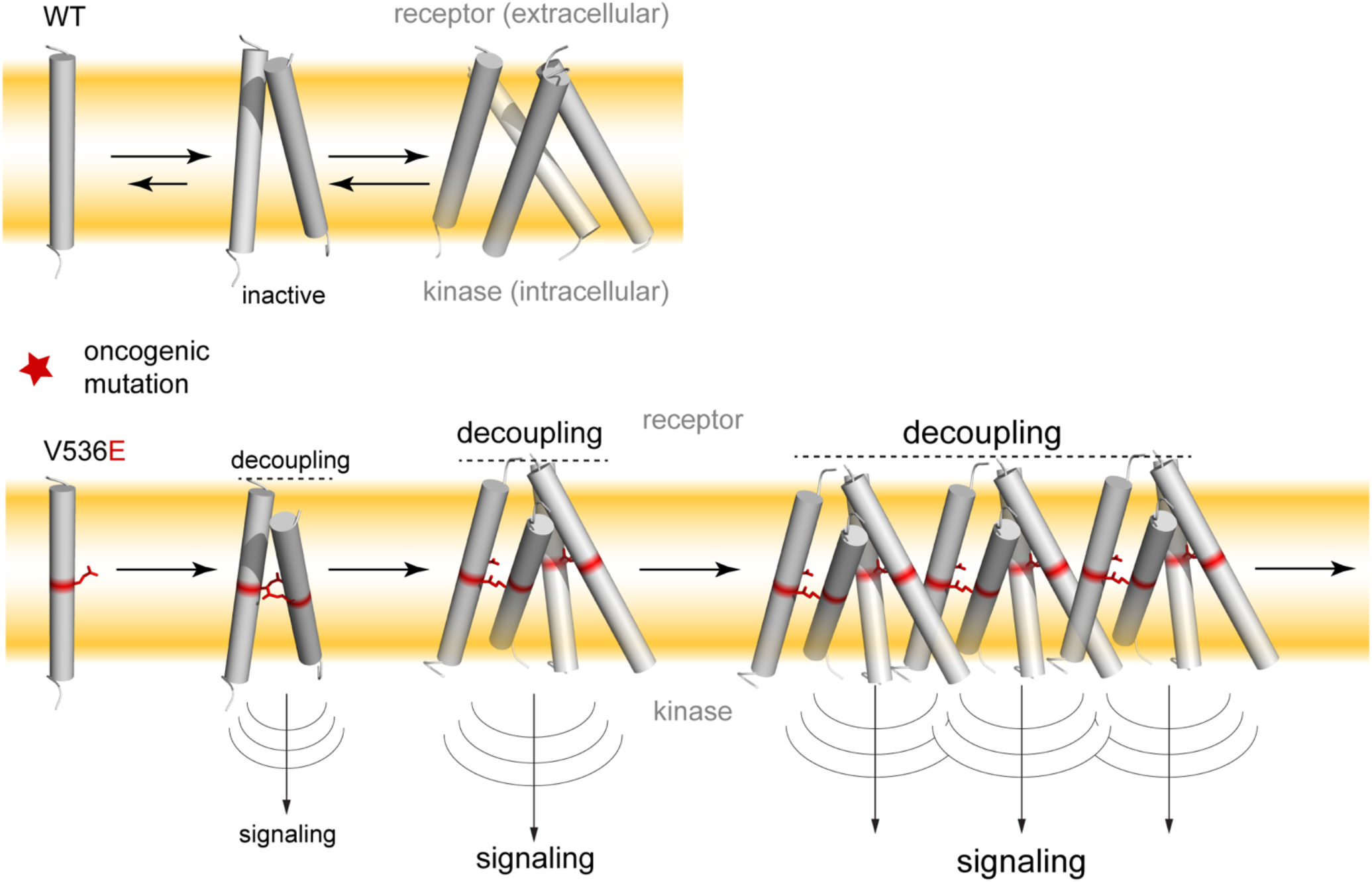
Hypothetical scheme of PDGFRA activation *via* TM oligomerization. WT TM domains have low tendency for dimer formation and further oligomerization is impaired (*upper panel*). Oncogenic mutation (e. g. V536E, shown with red sticks) promotes TM domain oligomerization and subsequent activation, whereby tetramer represents a building block in chain oligomerization (*lower panel*).

Our results give a novel perspective for potential biomedical applications and can be used as a starting point in development of new drugs and therapeutic strategies to control PDGFRA functioning and to combat pathologies caused by its dysfunction. At the same time, we should highlight an important fundamental aspect of the presented data and conclusions. In the particular example of PDGFRA, it was shown for the first time that for single-pass membrane proteins (RTKs, etc.), knowledge of the molecular details of their TM dimerization may not be sufficient to understand the cell functioning of the receptor(s), if this is associated with higher-order oligomerization, e. g. with the formation of tetramers. Thus, within the presented mechanistic framework it is shown that the structure and dynamics of the latter can radically differ from the properties of their two constituent dimers pointing toward synergy in behavior of the system upon changing in its oligomeric state. In addition, a comparative analysis of TM domains in the WT and in the V536E mutant revealed a fundamental difference in the behavior of the system as a whole, mainly due to drastic changes in corresponding dynamics of TM helices. Such effects are hard to detect only using modern instrumental methods, but based on *in silico* prediction, experiments can be planned to create and test new mutant forms of RTK with specified properties.

It is important to note that a number of additional factors also contributing to the oligomerization of PDGFRA are beyond the scope of the present study, where the receptor TM domains are mainly in focus. Thus, consideration of juxta membrane regions, receptor or/and kinase domains and at best - the full-length receptors for the future modeling will give better understanding of a possible structural organization of PDGFRA oligomers and realization of the receptor functional dynamics in that context. While for EGFR all-atom MD simulations of full-length receptor dimers (Arkhipov et al., 2013) or oligomers of their receptor parts (Zanetti-Domingues et al., 2018) have already been performed, in a case of PDGFRA this is still have to be carefully explored and also represents the current topic of our studies. At the same time, some important aspects of the structural and dynamic behavior of TM domains of RTKs were not addressed in the aforementioned studies. Also, the effects of pathogenic point mutations were not considered in a comparison to WT proteins. Another aspect is a potential role of membrane environment in the receptor oligomerization. As it was shown in a case of PDGFRB (Seong et al., 2017) and other receptors that their clustering and activation can be controlled by localization in the specific membrane environment (e.g. lipid rafts) (Bocharov, Mineev, et al., 2017) (Roy & Patra, 2022).

The proposed mechanism of PDGFRA constitutive activation via mutation-induced chain oligomerization gives an interesting prospective in understanding of the initial step of RTK related disorders. While being hypothetical it requires direct experimental validation *in vivo*, which may be laborious due to limited resolution and a number of known experimental biases (e. g., receptor overexpression typical for such experiments). At the same time, the oligomerization driven development of pathological states due to a single-point mutation in the PDGFRA TM domain relates this process to other known phenomena of this kind e.g., neurogenerative deceases whereby single-point mutations often lead to pathological protein aggregation (Ross & Poirier, 2004) (Bertram & Tanzi, 2005) (Vance et al., 2009) (Nomura et al., 2014). Interestingly, it was recently shown that human disease-related mutations also more likely increase protein aggregation than the diseases-unrelated ones and suggested that disease-associated protein aggregation represents rather widespread phenomena (De Baets, Van Doorn, Rousseau, & Schymkowitz, 2015). Thus, putative mechanism of PDGFRA activation *via* the oligomerization prone TM mutation may represent a realization of a universal principle in background of different human pathologies.

## 5. Conclusion

In this study we investigate a potential RTK activation mechanism due to TM mutations associated with higher-order receptor oligomerization. Using predicted configuration of TM tetramer for PDGFRA we demonstrate that the V536E mutant tetramer retains stable configuration on the microsecond timescale of MD simulations, while WT tetramer displays a clear tendency to dissociate. Apart from the tetramer stability, V536E alternates dynamics of TM helices within the tetrameric arrangement. While WT helices within the tetramer retain a piston-like motions en bloc that at the same time restricts lateral displacement of the C-termini and their dynamics, the mutant helices demonstrate increased dynamics in the C-terminal parts, which is also decoupled from more constrained N-termini. One of new interesting result of this work is that the structure and dynamics of RTKs’ tetramers can be significantly different from those of their dimers. Such a synergy may play an important functional role in cell signaling. The results of atomistic modeling open up a possibility to highlight a potential impact of higher-order receptor oligomerization induced by TM mutation into ligand-independent signaling of PDGFRA that may also be relevant to other members of RTK family. This a potentially new layer in the receptor constitutive activation may stimulate search for novel biomedical applications. While targeting TM domains of single-pass membrane receptor and modulating their dimerization and subsequently activation represents a promising therapeutic concept (e. g., “interceptor” peptides (Najumudeen, 2010) (Albrecht et al., 2020) (Westerfield & Barrera, 2020)), the putative activation scenario proposed here for RTKs invokes development of medical strategies to specifically inhibit and prevent pathological oligomerization in the TM region.

## Acknowledgements

The work was supported by the Russian Science Foundation (grant 18-14-00375 for RGE). Supercomputer calculations were performed within the framework of the HSE University Basic Research Program. Access to computational facilities of the Supercomputer Center “Polytechnical” at the St. Petersburg Polytechnic University is gratefully appreciated.

## Supplementary data

Supplementary Figure S1 and Movie S1, S2. Supplementary data to this article can be found online

## Supplementary Movie captions

**Movie S1**. Characteristic motions of TM helices in the WT tetramer. 1 µs MD trajectory filtered along the first three eigenvectors of the mass-weighted MD covariance matrix. Movie frame rate is 25 per second, whereby each frame is separated by 4 ns (0.1 µs per second). Helices are shown with cartoon representation.

**Movie S2**. Characteristic motions of TM helices in the V536E tetramer. E536 “hinge” residues are shown with sticks. Other details are given in the caption for Movie S1.

## Supplementary Figure caption

**Figure S1.**
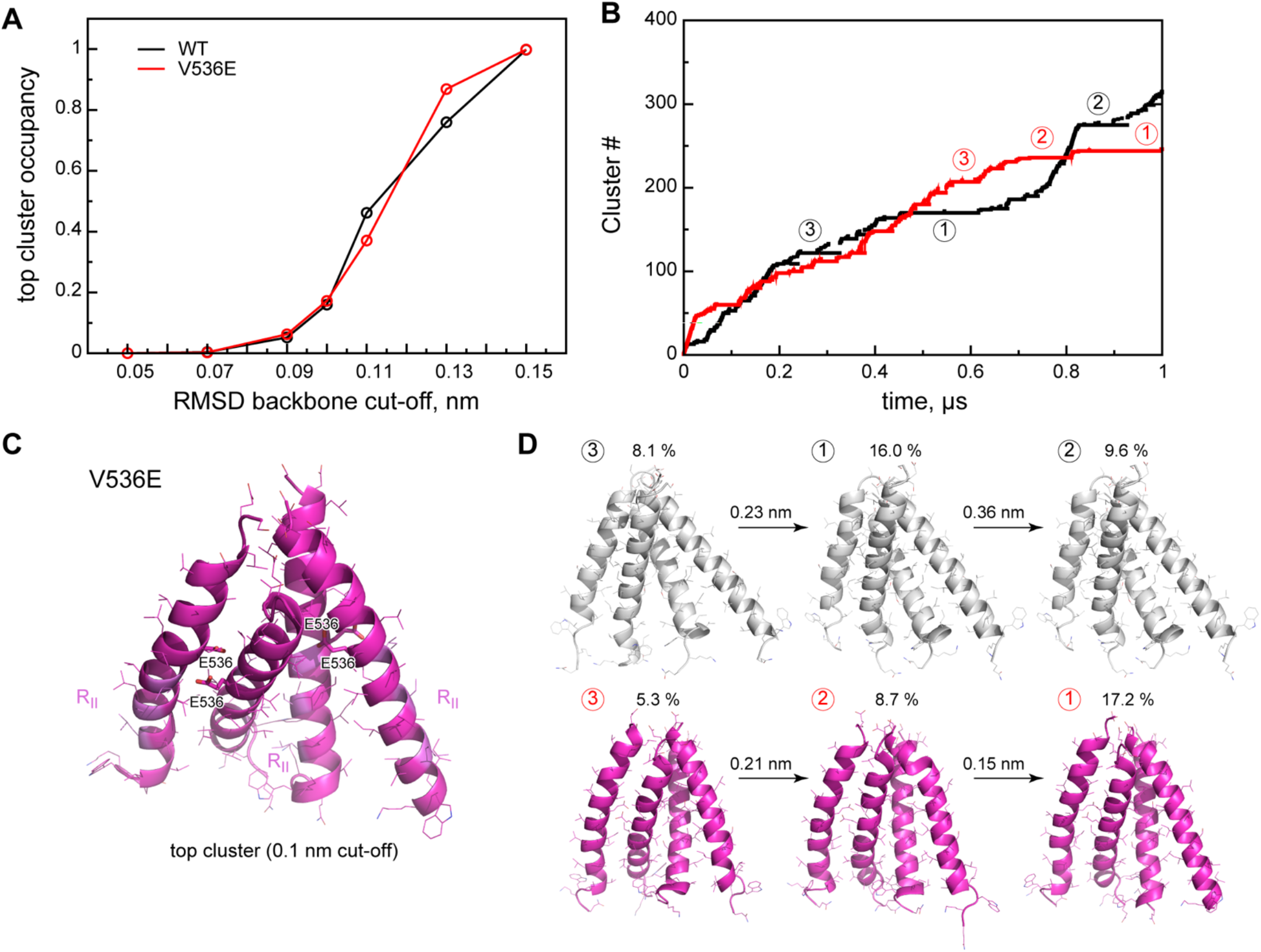
Cluster analysis of tetramer conformations. **A**. Distributions of the top (the most populated) conformation cluster occupancy as a function of backbone RMSD cut-off for WT (black) and V536E (red) tetramers. **B**. Conformational clusters along the MD simulation time (for the RMSD cut-off value of 0.1 nm). Positions of the first three most populated clusters are indicated with encircled numbers. The color code is the same as in panel A. **C**. The converged and the most populated conformational state (the RMSD cut-off value for clustering is 0.1 nm) of V53E tetramer shown in cartoon representation. Sidechains are shown with lines. E536 residues are shown with sticks and labeled. Oligomerization interface (R_II_) regions are shown with light magenta. **D**. Visualization of the top conformational clusters (see panel B) for WT (gray) and V535E tetramers given in cartoon representation with sidechains shown with lines. Cluster occupancy is indicated with percentages values above the structures. Corresponding RMSD values between the states are displayed above the arrows.

